# Does breeding season variation affect evolution of a sexual signaling trait in a tropical lizard clade?

**DOI:** 10.1101/864132

**Authors:** Levi N. Gray, Anthony J. Barley, David M. Hillis, Carlos J. Pavón-Vázquez, Steven Poe, Brittney A. White

## Abstract

Sexually selected traits can be expected to increase in importance when the period of sexual behavior is constrained, such as in seasonally restricted breeders. *Anolis* lizard male dewlaps are classic examples of multifaceted signaling traits, with demonstrated reproductive function reflected in courtship behavior. Fitch and Hillis found a correlation between dewlap size and seasonality in mainland *Anolis* using traditional statistical methods. Here, we present two tests of the Fitch-Hillis Hypothesis using new phylogenetic and morphological data sets for 44 species of Mexican *Anolis*. A significant relationship between dewlap size and seasonality is evident in phylogenetically uncorrected analyses but erodes once phylogeny is accounted for. This loss of statistical support for a relationship between a key aspect of dewlap morphology and seasonality also occurs within a species complex (*A. sericeus* group) that inhabits seasonal and aseasonal environments. Our results fail to support seasonality as a strong driver of evolution of *Anolis* dewlap size. We discuss the implications of our results and the difficulty of disentangling the strength of single mechanisms on trait evolution when multiple selection pressures are likely at play.

## Introduction

Signaling traits such as nuptial color in fish and bird songs can greatly affect survival and fitness of individuals, and thus can be used to understand how ecological and sexual selection drive phenotypic evolution (Endler & Basolo 1998; Cornwallis & Uller 2010). Postulated mechanisms for the evolution of these traits span numerous hypotheses of natural and sexual selection. Since many speciose radiations exhibit signaling traits, these traits are also regularly identified as “key innovations” (Panhuis et al. 2001). Despite their apparent importance for evolution, determining selective drivers for signaling traits has proven difficult (Cornwallis & Uller 2010).

Determining common mechanisms for the evolution of signaling traits is of great interest to biologists due in part to the vast interactions these traits initiate in nature. Signal trait evolution can be influenced not only by conspecifics via sexual selection (Darwin 1871; Kirkpatrick 1982; Ryan et al. 1993), but also by competitors (Rand & Williams 1970; Grether et al. 2009) and predators (Endler 1978; Tuttle & Ryan 1982) through natural selection processes. This multitude of processes therefore has the potential for downstream ecological and evolutionary effects on organisms and ecosystems. For instance, hypotheses driving the variation of a trait might provide key insights to patterns of diversification (sensory drive; Endler 1992) or community assembly (species recognition; Rand & Williams 1970). Mechanisms driving the evolution of sexually selected traits are important components of understanding biological processes on multiple scales.

*Anolis* lizards are common research subjects in evolution (Losos 2009). Among *Anolis* traits, dewlaps are perhaps the most discussed and least understood. Males of almost all ~400 anole species have dewlaps, which are flaps of gular skin used for species recognition, territorial behaviors, predator deterrence, and courtship (Losos 2009). Dewlap variation—most notably in size, color, and display characteristics—is considerable in *Anolis*, and studies have characterized general patterns across species (Fitch & Hillis 1984; Losos & Chu 1998; Nicholson et al. 2007; Ingram et al. 2016) and within species complexes (Ng et al. 2013, Driessens et al. 2017). To date, attempts incorporating many species have largely failed to find strong support for hypotheses to explain the evolution of dewlap diversity (Losos & Chu 1998; Nicholson et al. 2007; Ingram et al. 2016).

Mechanisms suggested to play important roles in creating dewlap variation include species recognition (Rand & Williams 1970), sensory drive (Fleishman 1992), and sexual selection (Fitch & Hillis 1984). The species recognition hypothesis suggests that dewlap variation evolved as a means for recognition of conspecifics (Losos 2009). This hypothesis garners support from the observation that many anole assemblages include multiple species with dewlaps that tend to vary greatly in size, pattern, or color (Rand & Williams 1970; Losos 2009). Quantitative support for species recognition driving evolution of dewlap traits across many species, however, has been elusive. Losos & Chu (1998) and Nicholson et al. (2007) tested for dewlap correlation with environmental, lineage, and assemblage factors across *Anolis* and found weak (nonsignificant; Losos & Chu 1998) support for sensory drive and no support for other hypotheses.

Sexual selection is the only hypothesis that has been supported in driving dewlap size across a broad sample of anoles (Fitch & Hillis 1984). Fitch & Hillis (1984) tested whether dewlap size correlates with length of the breeding season. Anole species living in seasonal areas were found to have shortened breeding seasons, while those in more aseasonal environments were found to breed throughout the year (Fitch 1972; Fleming & Hooker 1973; Andrews & Rand 1974). The authors suggested that species with short breeding seasons experience intense sexual selection relative to species that occupy aseasonal environments and breed potentially continuously (Fitch & Hillis 1984). Using data for 37 species, they found that anoles in seasonal environments had larger dewlaps than aseasonal species (Fitch & Hillis 1984). Species with larger dewlaps also exhibited stronger male-biased sexual size dimorphism, providing additional support for their hypothesis. *Anolis sericeus* (now considered a species complex; Lara-Tufiño et al 2016; Gray et al. 2019), the one species found in both seasonal and aseasonal habitats, fit their interspecies pattern: populations they sampled from seasonal environments possessed larger dewlaps than those from aseasonal environments. Their findings of seasonality environment affecting the evolution of a sexual signal could have important implications for sexual species that experience environmentally-imposed restrictions on length of the breeding season. A similar effect of length of breeding season on a sexual trait has since been documented in harvestmen (Burns et al. 2013), but to our knowledge has not been investigated in other systems.

An important advancement for testing hypotheses in evolutionary biology occurred with the development of methods for phylogenetic correction (Felsenstein 1985; Harvey & Pagel 1991). If dewlap size variation violates assumptions of independence among anole species, evolutionary history should be taken into account before concluding support for the Fitch-Hillis Hypothesis. Based on their sampling, there is evidence phylogenetic history could be a confounding factor in analyses (Fitch & Hillis 1984). For instance, most of the “seasonal” species (9 out of 16 species) sampled were from a single west Mexican clade that exhibits large dewlaps and occurs exclusively in seasonal areas (Poe et al. 2017). With a well-sampled and strongly-supported phylogeny (Poe et al. 2017), we can better address evolutionary questions via comparative methods that can account for phylogenetic non-independence of traits like dewlap size.

Here, we aim to assess support for the Fitch-Hillis Hypothesis of temporal constraint driving evolution of a sexual signaling trait. Specifically, we test whether species experiencing short breeding periods have relatively larger male dewlaps than those that can breed for more extended temporal periods. We test this contention at two scales. First, we test for a relationship between seasonality and dewlap size across Mexican anole species using phylogenetic regression. Second, we test for this relationship within silky anoles (*Anolis sericeus* complex), the only species group that occurs throughout highly seasonal and aseasonal environments in the region.

## Methods

### Data collection and measurements

We took digital photographs of individuals collected between 2010 and 2018 throughout Mexico, spanning all habitat types inhabited by anoles and some representing noteworthy distribution records. We only included images of adult males aligned flat with a calibration for proper measurement. As a proxy for body size, we used head length (HL; Ingram et al. 2016). Measurements for HL (in mm) and dewlap area (mm^2^) were taken using ImageJ (Schneider et al. 2012). Geographic coordinates were taken at collection sites.

In the original study, Fitch & Hillis used habitat type to categorize seasonal (desert, thorn-scrub, deciduous forest, and dry coniferous forest) versus aseasonal (tropical rainforest and cloud forest) environments in terms of rainfall (1984). Instead, we treated seasonality as a continuous variable. We extracted seasonality data from the seasonality of precipitation (BIO15) layer from WORLDCLIM (Hijmans et al. 2005) for each collection site using QGIS (QGIS Development Team 2016). These values represent the coefficient of variation of precipitation levels and should tightly correlate with length of breeding season for anoles (discussed further in Supplementary Material; Fig. 1).

**Figure 1:**
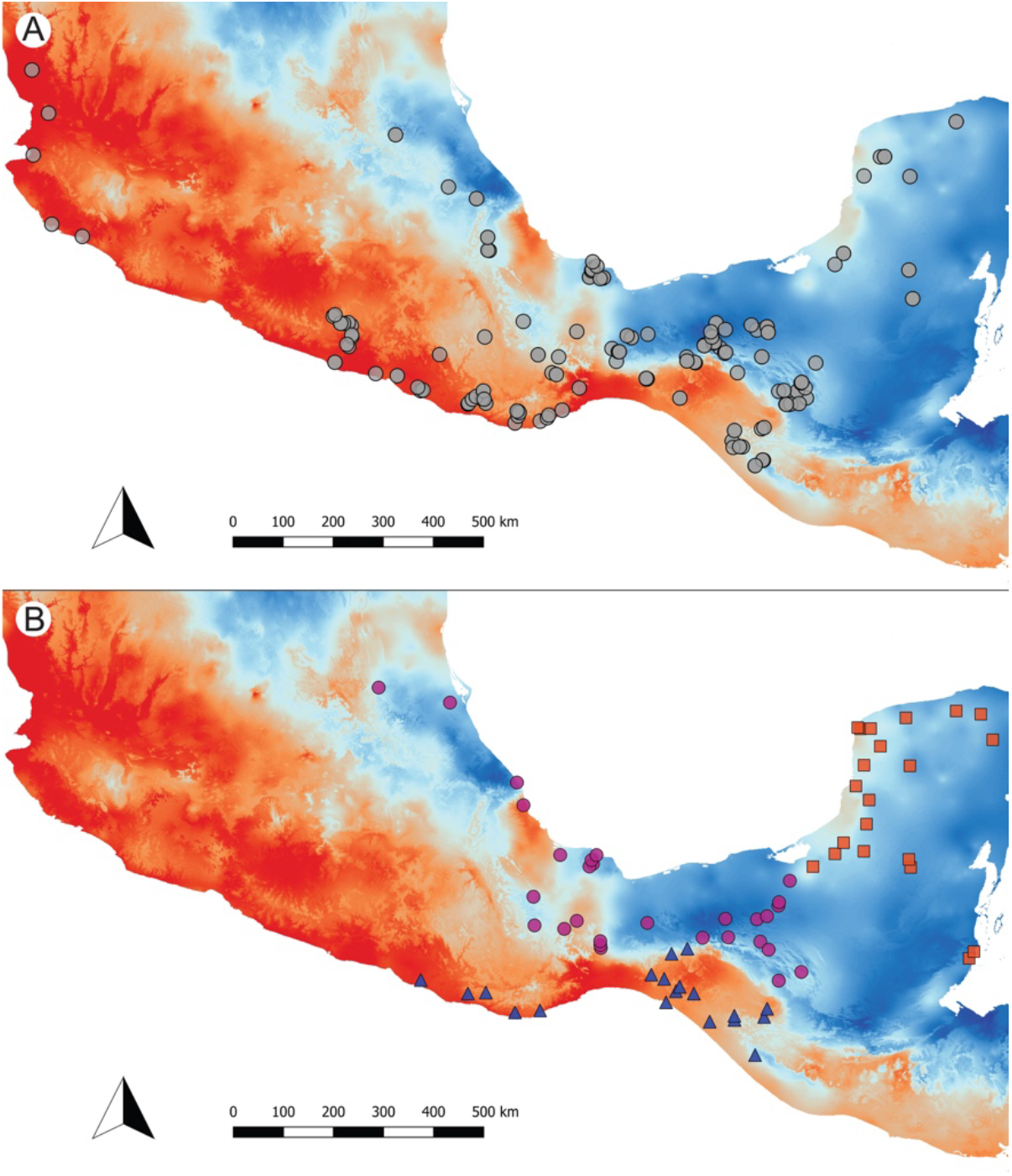
(A) Map of sampling localities with background raster reflecting seasonality (BIO15), with warmer colors reflecting high seasonality and cooler colors reflecting low seasonality. (B) Silky anole sampling, colored by clade. The Pacific clade (blue) occurs in the Pacific versant of southern Mexico, the Caribbean clade (purple) occurs south and west of the Yucatan in the Caribbean versant of Mexico, and the Yucatan clade (red) is only in the Yucatan Peninsula. These three lineages form a clade excluding the rest of the silky anoles, which occur outside Mexico.

### Species comparisons

Mexican anole taxonomy has a long history of uncertainty (Lieb 2001; Nieto-Montes de Oca et al. 2013). We analyzed all Mexican species for which we had data, and for which previous research supports their validity (n = 44; Tables S1, S2). To account for phylogeny, we used a recently published phylogeny (the maximum clade credibility tree for anoles from Poe et al. 2017), trimming the tips to match our sampled species using the ‘ape’ package in R (Paradis et al. 2004).

For each species, we averaged dewlap size and seasonality values from sampled localities. To verify consistency in both dewlap size and seasonality values within species, we calculated the intraclass correlation coefficient (ICC) for species for which we had at least 5 samples using the “irr” package in R. We calculated ICCs by randomly selecting 5 samples for each species before running analyses. Because body size is a confounding variable for dewlap size (Losos & Chu 1998), we ran a phylogenetic regression using the “ape” and “nlme” packages in R (R Core Team 2014) on log-transformed dewlap size with log-transformed HL for all species (Schoener 1970). We used the residual values from that regression as relative dewlap size for each species. We then performed phylogenetic regression on relative dewlap size against seasonality. To assess the strength of phylogenetic signal of BIO15, we calculated Pagel’s λ and Blomberg’s K. We also performed these analyses using standard (i.e., phylogenetically uncorrected) Ordinary Least Squares regressions.

In Mexico, the *Anolis sericeus* group (silky anoles; considered by Fitch & Hillis [1984] to be a single species) consists of three divergent clades (referred to here as Pacific, Caribbean, and Yucatan) that may be separate species (Gray et al. 2019; Fig. 1, Supplementary Material). A phylogenetic regression was not possible for these analyses, as we sampled from a number of unsampled populations from Gray et al. (2019). We averaged dewlap size for specimens from each locality and performed a standard OLS regression of dewlap size and head length using all silky anole localities. Using residuals from that regression as measures for relative dewlap size, we then regressed those values against seasonality. Subsequently, we performed another regression analysis using localities from only the Pacific and Caribbean clades. The Yucatan population is diagnosed by small male dewlaps (Lara-Tufiño et al. 2016). By removing these forms, we tested whether the Yucatan is strongly influencing the results of the analysis when the group is analyzed as a whole. We also ran each lineage on its own to test for signal within each group.

We performed a series of alternative analyses to explore consistency of our results and test support for the Fitch-Hillis Hypothesis. We ran analyses using HL as a covariate in the regression, using absolute values of dewlap size (largest sample for each species and average dewlap size), and testing season of collection as a covariate. Details for these analyses are in the online supplement.

## Results

We compiled data for 230 adult male dewlaps representing 41 species for interspecies analyses. Sampling within species ranged from 1-38 individuals from 1-22 localities. Within-species variation in dewlap size and seasonality was minimal (Table S1), reflected in ICC scores of 0.768 in dewlap size (n=14 species; p<0.001) and 0.824 in seasonality (n=7 species; p<0.001). Data files, including those used in alternate analyses, are available on Dryad.

Our standard OLS regression resulted in a significant positive relationship between relative dewlap size and seasonality (p=0.01537; F-statistic=6.426, DF=39; Fig. 2), consistent with results from Fitch & Hillis (1984). Accounting for phylogenetic relationships rendered the relationship nonsignificant (p=0.5519; Fig. 2). Pagel’s λ was 1.089 and Blomberg’s K for BIO15 was 1.039, indicating strong phylogenetic signal for seasonality. These results were in agreement with all alternate PGLS analyses in failing to find a significant relationship between dewlap size and seasonality (see Supplementary Material for more details on analyses and results).

**Figure 2:**
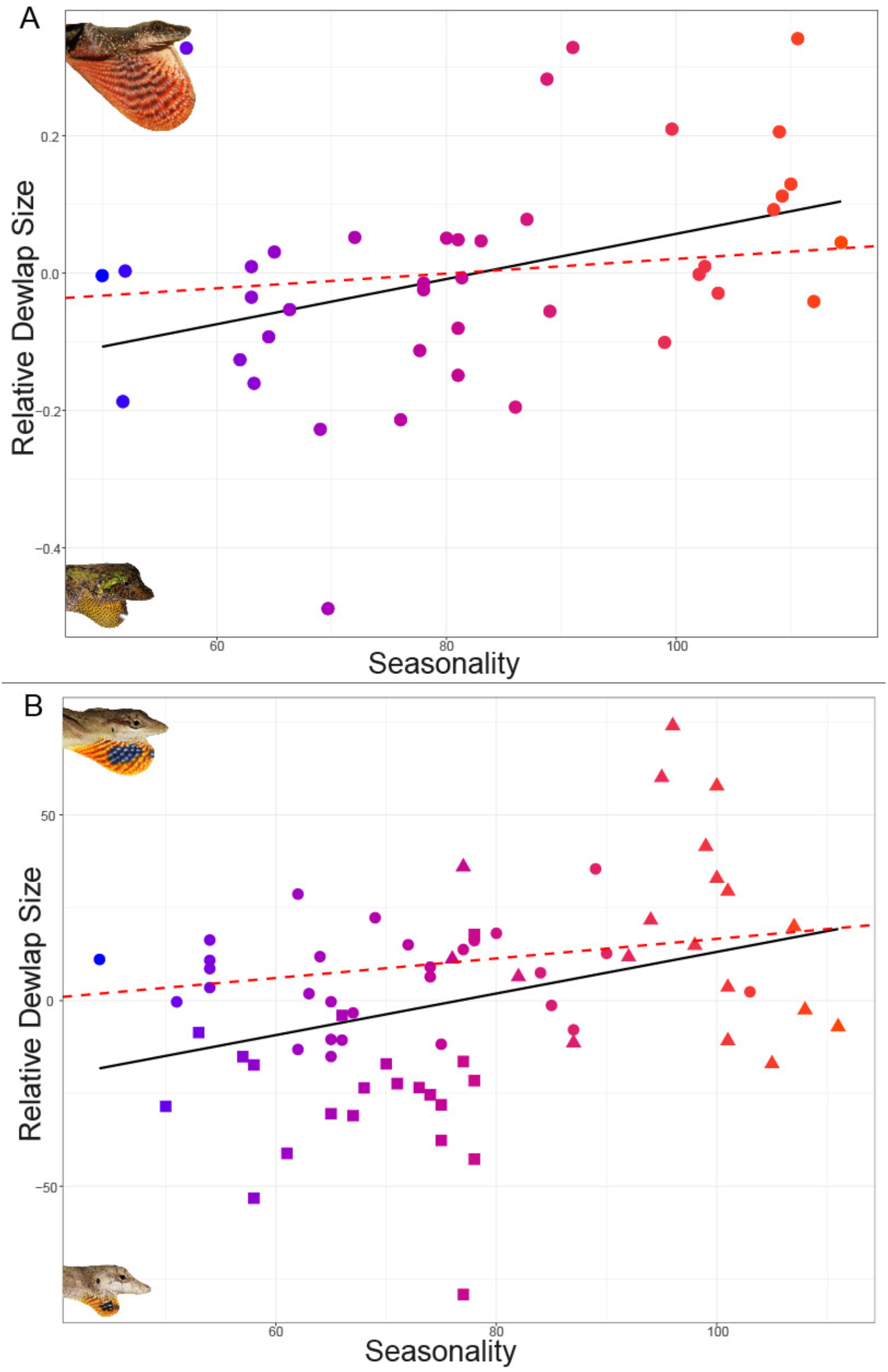
(A) Plot showing standard Ordinary Least Squares (OLS) regression (black line; p = 0.01537) and Phylogenetic Least Squares regression (dotted red line; p = 0.5519) of broad analyses. (B) Plot showing OLS regression of all silky anole populations (black line; p = 0.002742) and only Pacific and Caribbean silky anole populations (dotted red line; p = 0.1059). Plotted points are colored by seasonality, with warmer colors denoting stronger seasonality environments.

Sampling for the silky anoles included 153 adult males representing 69 populations/localities (19 Pacific, 29 Caribbean, and 21 Yucatan). OLS regression on the entire group produced a significant positive relationship between relative dewlap size and seasonality (p=0.002742, adjusted r^2^=0.1131, F-statistic=9.676, DF=67; Fig. 2). The regression incorporating only the Pacific and Caribbean lineages, however, is nonsignificant for relative dewlap size and seasonality (p=0.1059; Fig. 2). Each of the single lineage analyses resulted in nonsignificant results (Supplementary Material; Fig. S1).

## Discussion

In each of our analyses, the effect of seasonality on dewlap size was small enough that we could not confidently distinguish it from no effect once phylogeny was taken into account. Our results suggest that phylogenetic inertia is a reasonable alternative hypothesis for the pattern observed by Fitch & Hillis (1984). Results for the silky anole clade strongly depended on the exclusion or inclusion of the Yucatan lineage, whose individuals possess a small dewlap and inhabit a primarily aseasonal region. Individuals in the Caribbean lineage (sister to the Yucatan; Gray et al. 2019) do not have a reduced dewlap and occur in statistically indistinguishable seasonality environments (Table 1). If seasonality affects the evolution of dewlap size in silky anoles, we would expect to see reduced dewlaps in both lineages found in relatively aseasonal environments or a significant increase in dewlap size in the lineage inhabiting seasonal environments, neither of which occurs.

**Table 1:** Silky anole sampling with average and range of seasonality experienced by each lineage. The Caribbean and Yucatan lineages occur in indistinguishable seasonality environments (t-test, p = 0.5896).

| Lineage | Sample Number | Number of Localities | Average Seasonality | Range of Seasonality |
| --- | --- | --- | --- | --- |
| **Caribbean** | 63 | 29 | 69.9 | 44-103 |
| **Pacific** | 44 | 19 | 96.3 | 76-111 |
| **Yucatan** | 46 | 21 | 68 | 50-78 |

Our results and those of others (Losos & Chu 1998; Nicholson et al. 2007) suggest that sexual selection, species recognition, and sensory drive hypotheses are not supported as sole explanations driving dewlap size in large multispecies analyses of anoles. However, dewlaps vary along multiple axes, including color and display mechanics, which may jointly shape phenotypic evolution. Variability in dewlap color, pattern, size, and use is considerable in *Anolis*, and unlikely to be the result of a single selective force (Losos & Chu 1998). Experimental studies may disentangle causal mechanisms for dewlap evolution (Driessens et al. 2017, Leal & Fleishman 2003), though in-depth studies of the widespread species *A. sagrei* have had limited success in this endeavor (Baeckens et al. 2018). As additional data on other aspects of dewlap traits become available (color, UV reflectance, pattern, display characteristics, etc.), more complex tests of hypotheses may be feasible.

We note that while none of our analyses that take evolutionary history into account found a significant relationship between dewlap size and seasonality, our results do exhibit positive slopes (Fig. 2). These results could indicate a minor effect of seasonality on dewlap size that is too weak for our current methods and sampling to disentangle (we also found strong phylogenetic signal for BIO15, our proxy for seasonality). Though we found consistency in dewlap sizes within species, one species outside the tropics is known to exhibit plasticity in this trait (*Anolis carolinensis*; Lailvaux et al. 2015). This type of plasticity, if present in tropical anoles, might obscure a relationship between dewlap size and seasonality, particularly when within species sampling is low. However, plasticity in *A. carolinensis* is associated with skin elasticity and a lack of use during the winter when the lizards are inactive. Anoles in tropical regions are still active outside of the breeding season (Fleming & Hooker 1973; Henderson & Fitch 1975; Gray & White, pers. Obs.). We also found no effect of season of collection affecting dewlap size in our dataset when we tested for this explicitly (Supplementary Material), though the possibility of plasticity in tropical anoles awaits more thorough investigation.

Looking closely at patterns within taxonomic groups casts more doubt on the strength of any putative effect of the length of breeding season on dewlap size in Mexican anoles. Among species in the monophyletic west Mexican anole clade, which we sampled entirely in this study (Poe et al. 2017), dewlap sizes tend to be large and species tend to occur in seasonal environments. However, the species with the largest dewlap in the group, *Anolis macrinii*, also occurs in the least seasonal environment (Table S1, Supplementary Material). Another large-dewlapped species in our study, the semi-aquatic *A. barkeri*, occurs in some of the least seasonal environments of all Mexican anoles. Since other semi-aquatic species are also known to have large dewlaps in Central America (Gray et al. 2016; Ingram et al. 2016), it may be tempting to assume that the semi-aquatic lifestyle leads to large dewlaps. However, the Cuban species *A. vermiculatus* may be the most aquatic anole and is one of the only anole species that entirely lacks a dewlap. These observations demonstrate the need for more complete data sets to avoid drawing conclusions based on small sample sizes, and that perhaps a number of factors contribute to dewlap trait variation in anoles as expected for complex signaling traits (Endler 1992).

In this study, we use a larger data set and more rigorous methods to test the Fitch-Hillis Hypothesis in *Anolis* lizards (Fitch & Hillis 1984). If length of the breeding season does not explain evolution of dewlap size in anoles, will another hypothesis yield strong support across a broad sampling of anole species? We have our doubts, as mechanisms shaping the evolution of sexual traits are often difficult to determine in empirical systems (Cornwallis & Uller 2010). Additionally, the complexity of sexual trait evolution is such that testing a single mechanism is likely to fail (Endler 1992). We suspect demographic and natural history characteristics may explain some of the dewlap size variation observed among anole species. For instance, sensory drive is likely an important factor in some anole species (Leal & Fleishman 2003) but its strength as a driver for dewlap evolution is partly dependent on the likelihood that a particular female will encounter multiple males or will have a preference for males exhibiting larger or more visible dewlaps. One could also envision a scenario in which the opposite of the Fitch-Hillis Hypothesis is true: species/populations with shorter breeding seasons could yield weaker selection on male dewlap size due to limited time for females to seek out preferred males. Unfortunately, data on abundance and relevant natural history traits are lacking for the vast majority of anole species. Taxonomic groups without some of these complications or with more complete natural history data may be better suited for testing the Fitch-Hillis Hypothesis. Groups containing variation in length of breeding season and in signaling traits are common and can provide further resolution as to the efficacy of the hypothesis moving forward. Advances in understanding female choice since the Fitch & Hillis study (1984) have only strengthened the potential relevance of their hypothesis; females commonly “prefer traits of greater quantity” (Ryan & Keddy-Hector 1992; Andersson 1994). Examples of putative study subjects span a broad swath of animal life, including spiders (*Habronattus pugillis*, Maddison & McMahon 2000), insects (odonates, Serrano-Meneses et al. 2008), frogs (*Acris crepitans*, Ryan & Wilczynski 1991), birds (*Ficedula hypoleuca*, Järvi et al. 1987), and fish (Malawi cichlids, Marsh et al. 1986). The availability of natural history data is without question the biggest limitation in testing the Fitch-Hillis Hypothesis and other hypotheses addressing evolutionary processes in empirical systems.

Given our results in these analyses, there currently is limited support for temporal constraint driving evolution of sexual traits (but see Burns et al. 2013). Although Fitch & Hillis (1984) suggested dewlap color could be a mitigating factor for size in anoles, our preliminary data do not support this idea. Species with small dewlaps in seasonal environments do not have categorically “brighter” dewlaps than those with large dewlaps (Supplementary Material; Table S1) and incorporating color into future analyses will be challenging. Perhaps approaches focusing on additional factors (UV reflectance, display characteristics, etc.) and overall signal conspicuousness (Endler 1992) or additional groups of anoles distributed across seasonal and aseasonal environments will find stronger support for a modification of the Fitch-Hillis Hypothesis in anoles. We hope that our study will encourage further research in testing the effect of temporal constraint on sexually selected traits in other groups, as the Fitch-Hillis Hypothesis remains theoretically plausible and untested in a number of appropriate sexual signaling systems.

## Acknowledgments

We are grateful to the many field assistants that helped locate and/or photograph lizards for this study.

## Author contributions

LNG conceived the study. LNG, AJB, CJPV, BAW, and SP collected samples and took photos. LNG collected data and LNG, AJB, and BAW performed analyses. All authors wrote the manuscript and discussed result interpretation. All authors are accountable for the content and approve the final version of this manuscript.

## Ethics

Research approved by UNM IACUC (16-200554-MC).

## Data accessibility

Data have been deposited at Dryad and will be made available upon peer-reviewed acceptance.

