## Supplementary Material for "Does breeding season variation affect evolution of a sexual signaling trait in a tropical lizard clade?"

### Sampling and analyses

Our samples were collected primarily in the winter (late November-February) and summer (June-September). The lead author was involved in all dewlap photos, either in taking the photo or holding the animals. Dewlaps were extended by using forceps to pull the hyoid, and only dewlaps that were fully extended were used in this study. Grid backgrounds were included in every dewlap photo to scale measurements in all photos. The lead author outlined the dewlap for each photo and calculated the area on ImageJ after using the background grid for scale.

*Anolis* lizards tend to have restricted breeding seasons tied to patterns of precipitation (Fleming & Hooker 1973; Fitch & Hillis 1984; Losos 2009). The BIO15 variable from WORLDCLIM is based on precipitation data and is used as a proxy for the length of the breeding season in *Anolis* lizards. The variable is calculated as the coefficient of variation of the monthly precipitation (O'Donnell & Ignizio 2012). In effect, this variable reflects "evenness" of rainfall, and therefore is directly connected to the length in a year in which conditions are acceptable for *Anolis* lizard reproduction. High BIO15 values, as reflected in the Pacific portion of the Isthmus of Tehuantepec (Figure 1), are related to strong decreases in precipitation during the dry season when reproduction is unable to occur in anoles (Fleming & Hooker 1973). Exceptionally high values of BIO15 (above 100, commonly observed near the Pacific coast of southern and central Mexico) were investigated by O'Donnell & Ignizio (2012) and found to be regions where the variance of the precipitation "exceeded the average precipitation," and therefore represent good examples of environments where the breeding season is likely to be truncated and lead to the heightened levels of sexual selection discussed by Fitch & Hillis (1984). Lower values of BIO15 are seen in areas that tend to be consistently wet such as in lowland Caribbean rainforests and cloud forests (Figure 1) and therefore offer a longer period for reproduction, as described by Fitch and Hillis (1984; see also Fitch 1972).

### Taxonomic decisions

#### *Excluded taxa in broad species comparison*

We decided it would be premature to recognize several recently-described Mexican anole species that lacked strong evidence supporting their recognition because including taxa that do not properly reflect species-level diversity could potentially bias the results of our analyses. The populations we are not recognizing as separate species are not distinguishable by dewlap size and occur in the same seasonality environments as the species from which they were suggested to be split from.

We excluded taxa that were described primarily on the basis of differences in hemipenial morphology between populations (Köhler & Veseley 2010; Köhler et al. 2010; Köhler et al.

2014). The authors primary justification for describing these taxa (despite continuous distribution of populations) was that hemipenial traits should be associated with reproductive isolation. In the species groups that have been investigated with multiple lines of molecular and/or morphological evidence, this hypothesis has not been supported. For instance, there are now several documented cases of within-species and within-population variation in hemipenial morphology (Köhler et al. 2012; Phillips et al. 2015; Lara-Tufiño et al. 2016; Gray et al. 2019). Finding variation in hemipenial morphology within species is to be expected due to high rates of hemipenial evolution (Klaczko et al. 2015) and the resulting likelihood of multiple hemipenial forms to exist prior to speciation (as expected with other traits not associated with reproductive isolation; Lara-Tufiño et al. 2016). Gray et al. (2019) used a large genomic (restriction site-associated DNA; RAD) data set to infer phylogeographic relationships in the silky anoles (*Anolis sericeus* complex) and found no association between population divergence and hemipenial morphology. The two taxa described based on hemipenial morphology that we excluded from analyses, *A. carlliebi* and *A. sacamecatensis*, were poorly sampled throughout the group's continuous distribution, making it difficult to make a strong case in support of recognizing the putative species as valid at this point in time (Köhler et al. 2014).

We also excluded taxa that were described solely on the basis of varying levels of divergence among mitochondrial haplotypes (Köhler et al. 2014). There is an abundance of evidence that mtDNA haplotypes can exhibit strong patterns of divergence without barriers to gene flow (Irwin 2002; Funk & Omland 2003; Petit & Excoffier 2009), and anole systems have been known to show significant mitochondrial divergence between freely-interbreeding populations (Thorpe et al. 2008; Thorpe et al. 2010; Ng & Glor 2011; Ng et al. 2016). Anole populations delimited as separate species by Köhler et al. (2014) exhibit far lower levels of mtDNA divergence than other populations known to maintain evolutionary cohesion (Thorpe et. al. 2008; Ng et al. 2011). This study also lacked sampling from near putative contact zones of the populations they described as separate species, which causes further difficulty in interpreting their results. A standard isolation-by-distance model could potentially explain the resulting phylogenetic structure in their analyses, and in the absence of evidence for reduced or absent gene flow it would be questionable to assume the continuously distributed populations are on independent trajectories.

For further information on species not recognized in our study, see Table S2.

*Silky anole sampling*

We grouped populations by clades recovered via phylogenetic analysis of a large multilocus genomic data set (Gray et al. 2019) rather than species delimited using hemipenial morphology (Köhler & Vesely 2010). The molecular data used in that study consists of over 500 restriction-site associated DNA (RAD) markers and attained strong resolution for clades within the silky anole group. The populations present in Mexico represent a monophyletic group that is deeply divergent from populations in Guatemala, Honduras, and to the southeast. Only the Yucatan lineage has been shown to be morphologically distinct. The name *Anolis ustus* has been resurrected for this lineage, which can be diagnosed by a smaller dewlap in males (relative to other silky anoles) and a slightly larger dewlap in females (Lara-Tufiño et al. 2016).

**Alternate analyses**

We also tested for correlation between dewlap size and seasonality with head length (HL) as a covariate. The results for these additional analyses were similar to our analyses testing relative dewlap size versus seasonality (p=0.0147, adjusted R-squared=0.6821, F-statistic=43.91, 38 degrees of freedom).

Results of the phylogenetically corrected interspecies analysis using HL as a covariate were also similar to the analyses presented in the main text (p=0.5567). Pagel's Lambda model using log HL as a covariate resulted in a value of 1.089 (the same Lambda value from the analyses in the main body of the manuscript).

For the silky anole analyses, using HL as a covariate on the whole group resulted in significant statistical support (p=0.00657, adjusted r-squared=0.5321, F-statistic=37.39, 62 degrees of freedom). Removing the Yucatan lineage again resulted in a non-significant result: p=0.152 (Fstatistic=36.72; 44 degrees of freedom). None of the analyses of individual lineages yielded significant results (Caribbean, p=0.623; Pacific, p=0.5836; Yucatan, p=0.653).

Rather than relative dewlap size, it is possible that absolute dewlap size is more important for signal efficacy. Analyses using maximum absolute dewlap size (largest among our samples) on the interspecies data set yielded nonsignificant relationships between dewlap size and seasonality (OLS: p=0.9129, adjusted r-squared=-0.02532, F-statistic=0.01213, 39 degrees of freedom; PGLS: p=0.3676). Using average dewlap size rather than maximum attained the same result (OLS: p=0.6569, adjusted r-squared=-0.0204, F-statistic=0.2003, 39 degrees of freedom; PGLS: p=0.9494).

Lailvaux et al.'s (2015) study demonstrating seasonal plasticity in dewlap size in *Anolis carolinensis*, if widespread in the genus, could be a confounding factor in our analyses. Our ICC analyses indicate strong consistency in dewlap sizes within species (including individuals collected across multiple seasons, which suggests seasonal plasticity is not widespread in species in our data set).

In another attempt to verify whether seasonal plasticity could be driving the relationships, we ran analyses on the interspecies data set using season of collection as a covariate. We categorized season of collection as either Wet or Dry for each sample and pared down the data set so that only samples from the best-sampled season were used. Samples categorized as "Dry" season were from late November to February, which is within the period of low rainfall (Janzen 1967; Müller 1996) in tropical Mexico. We then ran OLS and PGLS on this data set using season of collection as covariates. The OLS results were marginally significant (p=0.04904, adjusted r-squared=0.1018, F-statistic=3.267), while the results of the PGLS analysis were nonsignificant (p=0.6049). Season was not found to be a significant factor in either OLS (p=0.3149) or PGLS analyses (p=0.2615).

**Table S1**

Table providing details on per species sampling, range in dewlap size, seasonality of sampled localities, and dewlap color. For dewlap color, up to three colors are listed: primary color, secondary color, and tertiary color are ranked by proportion of each color in the dewlap.

| *Anolis* Species | No. of Samples | No. of Localities | Range logdsize | Range Seasonality | Color |
|---|---|---|---|---|---|
| ***alvarezdeltoroi*** | 3 | 2 | 2.51-2.62 | 55-71 | Red |
| ***barkeri*** | 7 | 4 | 2.69-3.13 | 43-66 | Red/purple/white |
| ***beckeri*** | 8 | 5 | 1.68-2.36 | 44-54 | Pink |
| ***biporcatus*** | 2 | 2 | 2.62-2.76 | 64-74 | Blue/orange/white |
| ***boulengerianus*** | 11 | 9 | 1.94-2.5 | 92-115 | Orange/yellow |
| ***campbelli*** | 2 | 1 | 2.4 | 65 | Pink |
| ***capito*** | 3 | 3 | 2.01-2.22 | 68-71 | Yellow |
| ***compressicauda*** | 3 | 3 | 2.37-2.41 | 44-57 | Purple/yellow |
| ***crassulus*** | 4 | 3 | 2.13-2.47 | 75-85 | Orange |
| ***cristifer*** | 1 | 1 | 2.27 | 86 | Red |
| ***cuprinus*** | 4 | 1 | 2.17-2.41 | 87 | Purple |
| ***cymbops*** | 1 | 1 | 2.37 | 79 | Pink |
| ***dollfusianus*** | 3 | 2 | 2.05-2.07 | 75-81 | Yellow |
| ***duellmani*** | 7 | 3 | 1.97-2.37 | 63 | Purple |
| ***dunni*** | 2 | 2 | 2.29-2.48 | 106-111 | Red/yellow |
| ***gadovii*** | 1 | 1 | 2.78 | 108 | Purple/pink |
| ***hobartsmithi*** | 6 | 4 | 2.2-2.37 | 48-52 | Purple |
| ***laeviventris*** | 16 | 12 | 1.71-2.17 | 50-99 | White |
| ***lemurinus*** | 38 | 15 | 1.84-2.52 | 53-76 | Red/orange |
| ***liogaster*** | 2 | 2 | 2.21-2.52 | 102-103 | Purple |
| ***macrinii*** | 1 | 1 | 3.14 | 90 | Orange/white |
| ***matudai*** | 3 | 3 | 2.24-2.31 | 83-99 | Purple |
| ***megapholidotus*** | 2 | 2 | 1.94-2 | 110-111 | Pink |
| ***microlepidotus*** | 1 | 1 | 2.28 | 102 | Orange/yellow |
| ***milleri*** | 1 | 1 | 2 | 81 | Pink-purple |
| ***naufragus*** | 6 | 3 | 2.15-2.38 | 68-78 | Orange-red |
| ***nebuloides*** | 13 | 10 | 1.97-2.63 | 93-109 | Pink |

| | | | | | |
|---|---|---|---|---|---|
| ***nebulosus*** | 8 | 8 | 1.72-2.28 | 110-126 | Orange |
| ***omiltemanus*** | 4 | 3 | 2.04-2.29 | 102-104 | Orange |
| ***parvicirculatus*** | 2 | 2 | 2.38-2.41 | 80-83 | Red/orange |
| ***petersi*** | 4 | 2 | 2.14-2.66 | 63-88 | Red/black |
| ***peucephilus*** | 1 | 1 | 1.93 | 99 | Orange |
| ***quercorum*** | 5 | 3 | 2.16-2.47 | 87-90 | Pink |
| ***rodriguezii*** | 33 | 22 | 1.59-2.03 | 43-89 | Orange-yellow/red |
| ***rubiginosus*** | 1 | 1 | 2.04 | 81 | Pink |
| ***schiedii*** | 2 | 1 | 2.4-2.44 | 78 | Orange |
| ***serranoi*** | 3 | 3 | 2.51-2.61 | 75-88 | Red/black |
| ***subocularis*** | 6 | 6 | 2.46-2.87 | 107-113 | Pink/yellow |
| ***taylori*** | 2 | 1 | 2.64-2.69 | 111 | Red/mint |
| ***tropidonotus*** | 3 | 2 | 2.09-2.3 | 65 | Yellow/red |
| ***uniformis*** | 5 | 4 | 1.9-2.12 | 53-66 | Purple/pink |

| Species not recognized | Synonymous with | Evidence for validity | Evidence against |
|---|---|---|---|
| ***Anolis nietoi*** | *Anolis nebuloides* | mtDNA clustering | Phylogenetic trees not consistent among markers; morphology and continuous distribution of population does not suggest reproductive isolation |
| ***Anolis stevepoei*** | *Anolis nebuloides* | mtDNA clustering | Phylogenetic trees not consistent among markers; morphology and continuous distribution of population does not suggest reproductive isolation |
| ***Anolis zapotecorum*** | *Anolis nebuloides* | mtDNA clustering | Phylogenetic trees not consistent among markers; morphology and continuous distribution of population does not suggest reproductive isolation |
| ***Anolis carlliebi*** | *Anolis quercorum* | mtDNA clustering/hemipenial morphology | Morphology and only slight hemipenial differences. Molecular sampling missing many important localities. |
| ***Anolis sacamecatensis*** | *Anolis quercorum* | mtDNA clustering/hemipenial morphology | Morphology and only slight hemipenial differences. Molecular sampling missing many important localities. |
| ***Anolis immaculogularis*** | *Anolis subocularis* | Distinct in dewlap coloration from one species and in mtDNA divergence from another species | Phylogenetic trees show a clustering of a mix of *immaculogularis* and *subocularis*; the population described is not distinct in morphology or in mtDNA. |

**Table S2**

Table summarizing evidence for and against recently described species not recognized in this study.

**Figure S1**

Figure showing best-fit Ordinary Least Square regression lines for each silky anole lineage. Red squares represent the Yucatan lineage, purple circles represent the Caribbean lineage, and blue triangles represent the Pacific lineage. The Caribbean regression line shows a positive slope, unlike the other two groups, but with a p-value of 0.623 it remains nonsignificant.

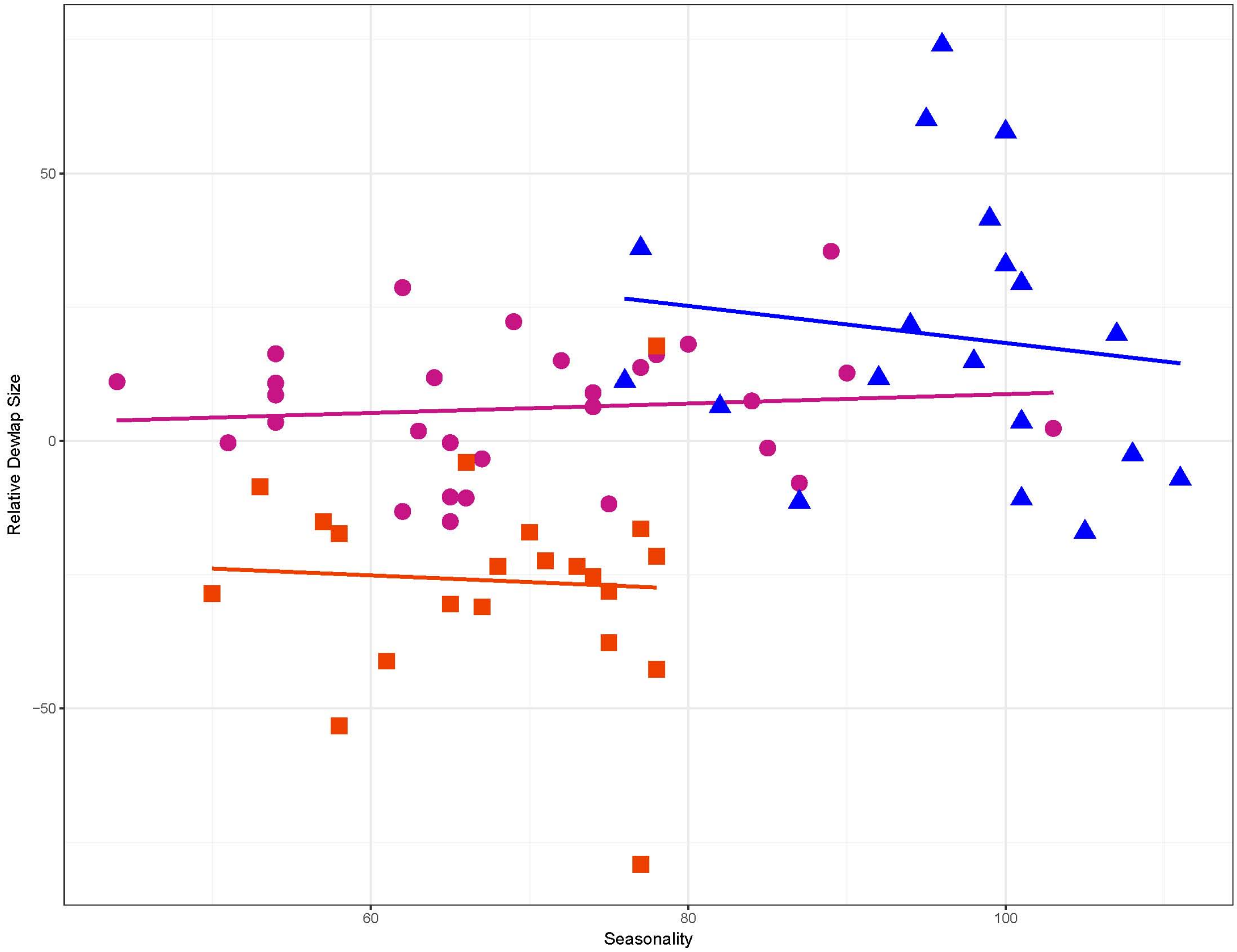
